# Explaining heritable variance in human character

**DOI:** 10.1101/446518

**Authors:** Jaime Derringer

**Affiliations:** Department of Psychology, University of Illinois at Urbana Champaign, Champaign, IL, USA

## Abstract

Personality is moderately heritable (around 40%), across a variety of measures [1]. Non-linear genetic influences (subsumed by the influence of the “dominance” factor in the heritability literature) likely play a large role in the genetics of personality [2], even more so than for other phenotypes. Zwir and colleagues [3] sought to identify sets of SNPs that, through both additive and interactive effects, explain the previously observed heritability of personality. The abstract reported identification of a set of SNPs that “explained nearly all the heritability expected for character in each sample (50 to 58%).” This reported effect size is extraordinary. The paper’s Supplemental Materials reveal how such an estimate occurred.

The variance explained in character by the identified SNPs are the R^2^ values from regression models in which polygenic scores were fit and evaluated within each of three samples (Finnish, N = 2 149; Korean, N = 1 052; and German, N = 902), without independent cross-validation either within or between samples [3]. This method as described, in which scoring weights are estimated and evaluated within a single sample without the use of cross-validation, is known to substantially inflate polygenic score effect sizes as a result of the model being overfit to the particular dataset [4]. These inflated estimates of variance explained within each sample were presented as evidence of strong aggregate effects of the identified genes that replicate across samples [3]. The reader may be reminded of the Sagan standard: “Extraordinary claims require extraordinary evidence.”

To initially reduce the number of SNPs for inclusion in subsequent analyses, Zwir and colleagues first identified SNPs meeting p < 0.01 in a GWAS of a single quantitative measure of character (a combination of three factors of Self-Directedness, Cooperativeness, and Self-Transcendence) within the Finnish sample. Next, they applied a data reduction technique to identify clusters among the retained SNPs, and tested the resulting 902 SNP sets for association with the single measure of character within the same Finnish sample. Forty-two of those SNP sets were identified as significantly associated with character. These 42 SNP sets included 2 971 unique SNPs [3] (Supplemental Table S3) representing 727 known genes.

These 2 971 SNPs were then brought forward to construct and evaluate a polygenic score within the same Finnish sample in which they had been identified. Here, no mention is made of cross-validation. The Supplemental Methods describe that the same scoring method was applied within each of the Korean and German samples, although the exact SNPs included in the polygenic scores differed between samples (indicating that the weights estimated in the Finnish sample were not applied across samples in a cross-validation sense; the exact weights applied were not reported) [3].

To put the claim of 50-58% of the variance in character explained by 2 971 SNPs into broader context, the schizophrenia polygenic scoring paper cited by the authors for the method applied identified a polygenic score that explained 3% of the validation-sample variance in schizophrenia (using independent discovery and validation samples of more than 3 000 participants each) [5]. To explore what might be a reasonable expected effect size for a personality polygenic score, Figure 1 summarizes results of 16 existing personality GWAS obtained from a central public database [6]. Across a variety of sample sizes and phenotypes, the SNP-based heritability estimates (which may be interpreted as the potential maximum effect size of a polygenic score) consistently fall below 12%.

**Figure 1.**
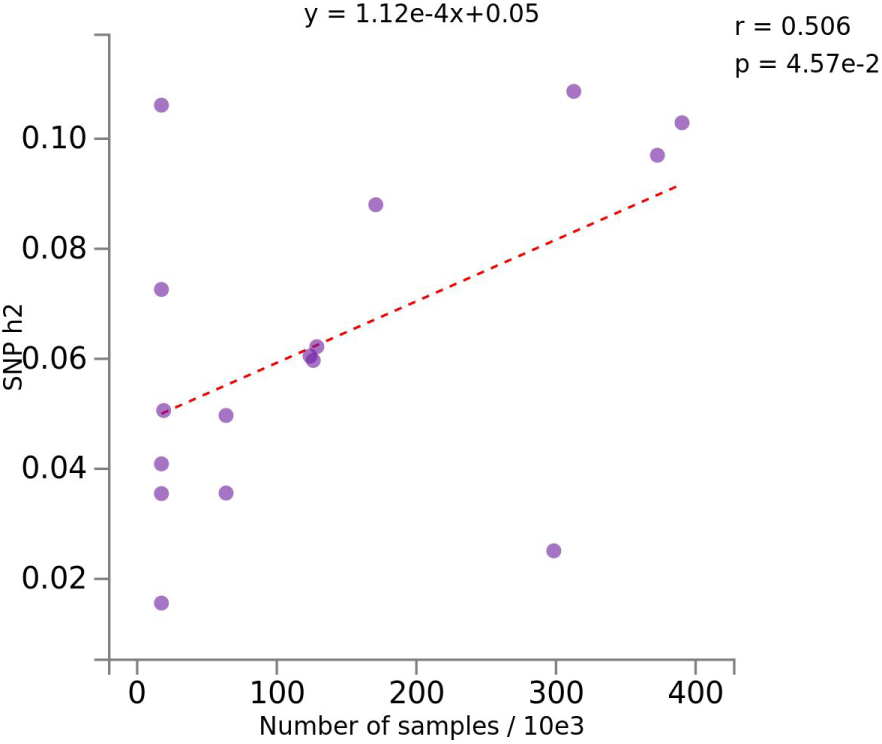
Sample size and estimated SNP heritability among personality GWAS.

Note: Figure generated using the Multiple GWAS Comparison tool from the GWAS Atlas [6]. GWAS of relevant personality traits were identified from among those whose summary statistics were available within the Psychiatric domain, Mental Functions chapter, Temperament and Personality subchapter, restricted to Big Five traits and those at a similar measurement level (e.g. disregarding symptom-level phenotypes) that represent face-valid construct overlap with the Character factors. The identified GWAS were [Atlas ID: trait name: PMID, if available, or ‘NA’ if unpublished results generated for the Atlas from UK Biobank data]: 30:Agreeableness:21173776, 26:Aggression:26087016, 3747:Life-Meaningfulness:NA, 31:Conscientiousness:21173776, 28:Extraversion:21173776, 33:Extraversion:26362575, 3385:Happiness:NA, 3745:Happiness:NA, 27:Neuroticism:21173776, 32:Neuroticism:24828478, 55:Neuroticism:27089181, 3417:Neuroticism:NA, 3795:Neuroticism:29942085, 29:Openness:21173776, 3296:Risk-Taking:NA, 54:Subjective-Well-Being:27089181

Note that Zwir and colleagues’ [3] target phenotype of character was established independent of any genetic information (Supplemental Methods, pp.4-5), so there is no reason to expect that it would be systematically more strongly heritable than other previously investigated arrangements of personality constructs. Further, the regression and resulting polygenic scores as described were the result of additive (main-effects-only) linear regression models (Supplemental Methods, p. 34), and so would not capture non-additive (interactive) influences more than any other model where additive influences are assumed (like SNP-based heritability). Polygenic scores have been consistently demonstrated to not substantially lose predictive validity with an increased number of included SNPs [5], so an additive polygenic score based on fewer SNPs, even if optimally identified, would not show a stronger effect size than a genome-wide additive polygenic score.

